# Mechanistic insights into state-dependent synchronization of neural activity by weak tACS fields

**DOI:** 10.64898/2026.11.20.768020

**Authors:** Eloise Habek, Joseph Tharayil, Weina Ji, James King, James B. Isbister, Michael W. Reimann, Georges Khazen, Niels Kuster, Taylor Newton, Esra Neufeld

## Abstract

Transcranial alternating current stimulation (tACS) biases spike timing with weak electric fields, but how re- current circuitry and network state shape this response is unclear. We simulated tACS in a detailed cortical column model in two network states of varying coherence, roughly corresponding to REM-like and NREM-like states. Despite weak impacts on network firing rates, stimulation entrained network activity at realistically occurring field strengths. Global entrainment was initiated by direct entrainment of ongoing pyramidal cell activity, which subsequently recruited inhibitory interneurons in a state-dependent manner. The NREM-like network locked more strongly to all tested stimulation frequencies, independent of baseline spectral power, and displayed larger recurrent-connectivity-mediated phase lags. However, recurrent connectivity was not required for, and did not amplify, entrainment, but instead produced an offset in baseline network activity. Our re- sults highlight the impact of network state on the response to tACS, and suggest that meso-scale models of electromagnetic-electrophysiological interactions must look beyond single-cell polarization and baseline oscilla- tions at the stimulation frequency.

## 1 Introduction

Transcranial alternating current stimulation (tACS) is a non-invasive subthreshold neuromodulation technique that interacts with ongoing brain rhythms to bias oscillatory spiking to a preferred phase with respect to the applied waveform, and which is under investigation both as a tool for modulating cognition in healthy users and as a candidate intervention in psychiatry and neurology [1, 2, 3]. Current experimental evidence suggests that tACS influences neuronal excitability, thereby entraining ongoing activity to the external stimulus, and in addition, can induce synaptic plasticity [2, 3, 4]. Here, entrainment refers to the emergence of a preferred phase relationship between spike times and the applied alternating current; this effect is also referred to as phase locking. *In vitro* recordings [5] and computational models of morphologically detailed neurons [6, 7, 8] demonstrate that the response to stimulation is cell-type-specific and depends on stimulus amplitude and frequency: phase locking increases approximately linearly with stimulus amplitude [6, 7], while dendritic asymmetry, cell length, and intrinsic oscillatory activity determine sensitivity to stimulation [7, 8], with large, slow-firing pyramidal cells responding strongly at low frequencies, and smaller, faster-firing inhibitory interneurons exhibiting peak responses at higher frequencies [5, 7, 8]. However, some inhibitory interneuron subtypes, particularly large basket cells, phase-lock to 10 Hz tACS comparably to layer 6 pyramidal cells [6].

While single-cell responses to weak electric fields are increasingly well understood, the mechanisms under- lying network responses to stimulation are less clear. *In vivo* recordings [9, 10, 11] and computational models [6] suggest that at field intensities typical of tACS in humans, spike timing, but not firing rate, is significantly influenced by stimulation. Computational and experimental studies have reported entrainment patterns con- sistent with an Arnold tongue: stimulation is most effective in entraining the network when applied at the frequency of an endogenous oscillation, with the range of stimulation frequencies that entrain an endogenous rhythm broadens as stimulation intensity increases; low-amplitude stimulation at other frequencies can dis- rupt endogenous activity [11, 12, 13]. The effects of brain stimulation have been found to depend strongly on network state, but the mechanisms by which this state dependence arises are unclear (see Bradley et al. [14] for a comprehensive review). For example, Rembado et al. [15], using intracortical sinusoidal stimulation in awake mice, found that spike-phase entrainment to stimulation, but not firing rate modulation, was strongly influenced by locomotor activity, and that this effect depended on stimulation frequency. However, a detailed account of the precise mechanisms at play remains elusive. While some studies [12] have highlighted the role of positive feedback through pyramidal-pyramidal connections in enhancing network responses to stimulation, others have suggested that pyramidal recruitment of inhibitory interneurons is primarily responsible for the observed synchronization of network activity [13]; regression analyses of brain-wide recordings, in turn, suggest that field-driven coupling rather than connectivity best predicts entrainment to low-frequency stimulation [15]. Surprisingly, while Rembado et al. [15] found that cells clustered with respect to their level of entrainment, these clusters did not demonstrate a clear cell-type preference, and might be more related to functional sub-networks.

Computational modeling of network activity in response to stimulation may shed light on the mechanisms of tACS, and advance its use in personalized medicine [16]. While several such models have been previously developed, most prior work has used either macroscopic and often phenomenological neural mass models or networks of point and two-compartment neurons. Recently, simplified spiking networks have been used to study the mechanisms underlying the intensity- and frequency-specific effects of tACS, which were shown to be attributable to network dynamics [17]. However, network models utilizing simplified neural morphologies often omit features that single-cell work identifies as decisive, such as dendritic morphology and cell-type- specific polarization; even point-neuron models that perfectly reproduce single-cell responses would be unable to capture phenomena such as electric field modulation of propagating activity in the neuronal arbor [18], which depend on the interplay between network state, morphology, and the applied field. Reduced models typically represent field effects through an effective input such as λ⃗ E⃗(t), where the sensitivity λ specifies the coupling between the applied field and neuronal input [19]; equivalent-input formulations may also incorporate frequency- dependent amplitude and phase filtering [20]. At the single-cell level, near-linear polarization with applied field strength is observed under subthreshold conditions [21, 22]. At the population level, however, λ becomes a semi- empirical parameter that aggregates morphology, orientation dispersion, synaptic state, and network context [19]. Even with constant coupling, nonlinear neuronal and network dynamics can produce amplitude-, frequency-, and state-dependent phase locking, so evaluating these formulations requires comparison with detailed circuit responses under matched stimulation and network conditions. While recent technological advances in mesoscale modeling [23, 24] have raised the possibility of predicting whole-brain responses to noninvasive brain stimulation, an insufficient understanding of the effects of stimulation at the microcircuit level means that such predictions are still limited by uncertainty in the coupling between the incident field and the mesoscale population model; whether a single, state-independent λ can reproduce the response of a cortical circuit to tACS is presently unknown.

Computational modeling at the microcircuit level may provide greater mechanistic understanding of the effects of noninvasive brain stimulation, and help identify parameters critically influencing the mesoscale re- sponse to stimulation. Previous work has examined tACS in a morphologically detailed, synaptically con- nected five-neuron microcircuit [25], with one representative neuron per cortical layer, which is too few to capture population-level dynamics or to represent a defined network state. Even in previous models of tACS in networks of point- and two-compartment neurons [12, 13, 17], synaptic connectivity, and especially, baseline network activity, have been substantially simplified, with models being tuned to produce specific oscillatory frequencies, thus limiting how readily any mechanistic insights generalize to more naturalistic brain states. While suprathreshold TMS has been studied in a large, detailed cortical circuit model [26], no such model has ever been applied to shed light on subthreshold electrical stimulation. Thus, whether and how weak alternating fields that only minimally polarize individual neurons influence recurrent networks with the cellular diversity and scale of a cortical column has not yet been studied, preventing bottom-up derivation and interpretation of realistic λ values.

To address this gap, we use the Blue Brain Project (BBP) model of rat non-barrel somatosensory cortex [27, 28], in which morphologically and electrophysiologically detailed neurons, realistic local connectivity, and spontaneous activity calibrated to *in vivo* firing rates coexist at the scale of a cortical column. We simulate a single column of approximately 30,000 neurons (Fig. 1a) and apply spatially uniform sinusoidal fields along its depth axis, parallel to the somato-dendritic axis of pyramidal cells—the orientation that produces the strongest somatic polarization in these cells [18, 21, 29, 30]. Stimulation is applied at three frequencies (7 Hz, 10 Hz and 15 Hz) and at subthreshold amplitudes from 0.25 V m*^−^*^1^ to 4 V m*^−^*^1^, in two network states that differ in synaptic efficacy and background drive. In particular, we simulate a network state with high correlation between excitatory and inhibitory populations, roughly corresponding to non-rapid eye movement sleep (hereafter referred to as the NREM-like condition), and a network state with low excitatory/inhibitory correlation, roughly corresponding to rapid eye movement sleep (hereafter referred to as the REM-like condition). It should be noted that the terms “REM-like” and “NREM-like” are primarily phenomenological descriptions of circuit activity rather than mechanistic explanations for that activity. However, there are biological parallels between the two model states and REM/NREM sleep. In particular, the reduced synaptic efficacy and increased variance in noise input in the model’s low-correlation state mirrors the effects of cholinergic neuromodulation in REM sleep; for further details see Sec. 3.3.1.

**Figure 1:**
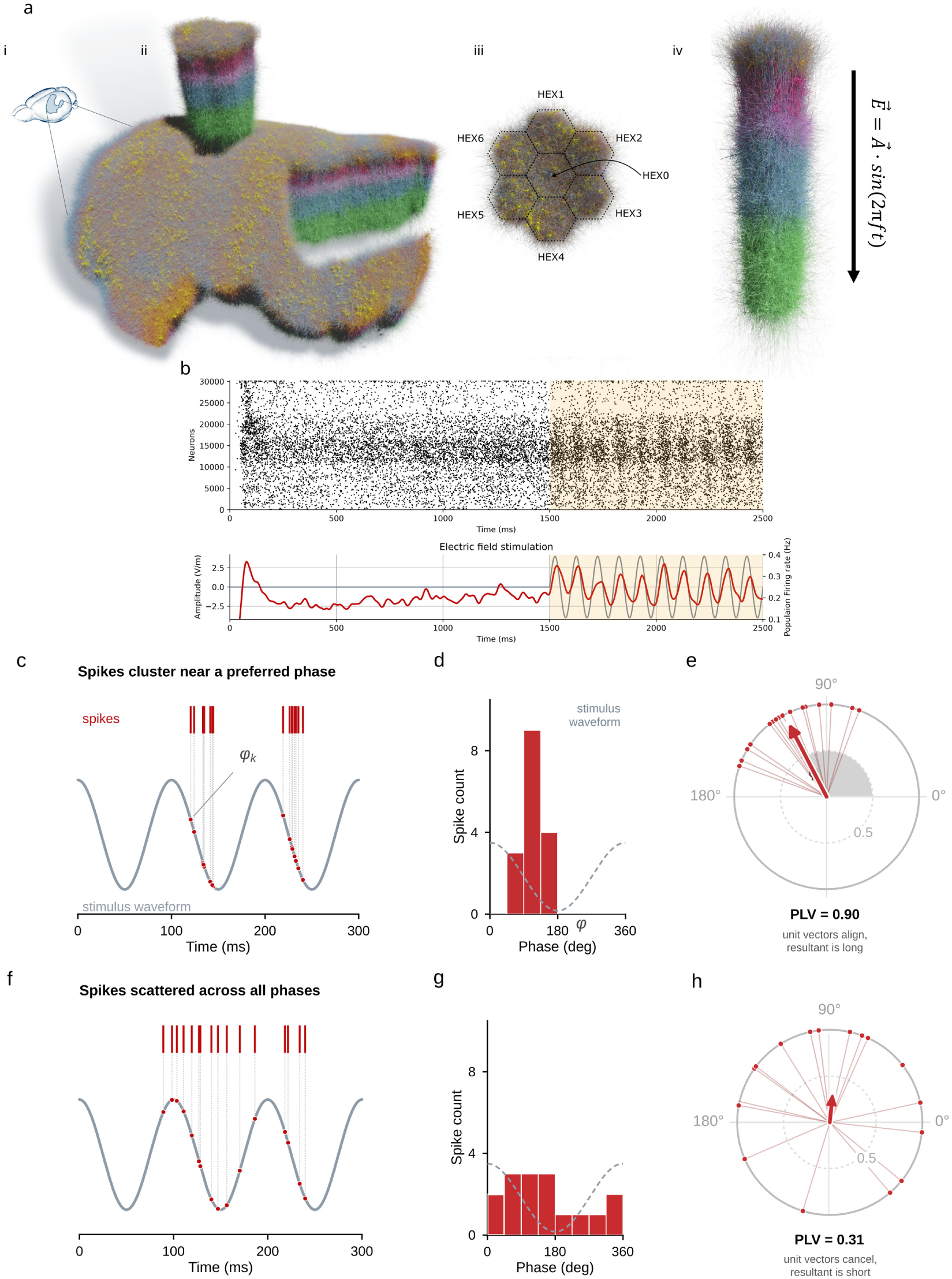
Model setup and example network response. a: Simulations are performed using either a seven- cortical-column subvolume (Supplementary Section S1) or a single-column subvolume (all other simulations) of a biophysically realistic model of the rat non-barrel somatosensory cortex developed by the Blue Brain Project. i: Non-barrel somatosensory cortex within the rat brain. ii: 7-column subvolume partially extruded from the somatosensory cortex. iii: Top-down view of the 7-column subvolume, highlighting the individual columns. iv: The single-column subvolume, featuring electrophysiologically and morphologically detailed neurons with 40 million synaptic connections with realistically-distributed strengths and locations. The applied field E(t) = A sin(2πft) is oriented along the depth axis of the column. b: Example simulation outputs for one stimulation condition (10 Hz, 4 V m*^−^*^1^ stimulus, REM-like state): raster plots of spiking activity (dots) and the stimulus waveform (gray) overlaid with the smoothed population firing rate (red). Stimulation begins at 1500 ms; the analysis window is the stimulation period from 1500 ms to 2500 ms (highlighted in yellow). c-e: The phase- locking value quantifies the consistency of spike timing relative to the stimulus. c: Spike times (ticks) from a simulated unit, shown against a 10 Hz stimulus waveform; dotted lines mark the phase ϕ*_k_* of the waveform at each spike. d: Distribution of the 16 spike phases; the dashed curve shows one cycle of the stimulus for orientation. e: Each spike contributes a unit vector e*^iϕk^* (thin lines); for spikes clustered near a preferred phase, the resultant (red arrow) has length PLV = 0.90, close to the maximum of 1. f-h: As in c-e, for the same number of spikes distributed approximately uniformly across phases; the unit vectors largely cancel and the resultant is short (PLV = 0.31). The PLV is bounded between 0 and 1 and is biased upward at low spike counts.

To separate the direct effect of the field from the contribution of stimulation-dependent recurrent feedback, we compare an intact circuit exposed to stimulation with a spike-replay condition. In the latter condition, the network is exposed to the same extracellular stimulus as the intact condition, but feedforward activity is blocked, and each neuron instead receives the synaptic input recorded from a simulation of a circuit with feedforward connectivity but without extracellular stimulation. The response of the spike-replay condition therefore isolates the direct impact of stimulation on the network, excluding effects mediated by recurrent connectivity. We quantify stimulus locking with the phase-locking value (PLV), which measures the concentration of spike phases relative to the applied waveform, and report the preferred phase separately (Fig. 1c-h); because the PLV is biased upward at low spike counts [31], we also report the pairwise phase consistency (PPC), a bias-free estimator of the squared PLV [32].

We show that spike-stimulus phase locking depends on field amplitude, frequency, and network state, in ways that are not predictable from single-cell models. Overall phase locking in the circuit increased nonlinearly with amplitude, with a gain that depended on stimulation frequency and network state. For the frequencies examined, the circuit response to stimulation is stronger in a network state with high correlation between excitatory and inhibitory populations (the NREM-like condition) than in a low correlation state (the REM- like condition); this difference is not attributable to differences in baseline oscillatory power at the stimulus frequencies. Moreover, we show that in the NREM-like state, recurrent connectivity plays a larger role in the network response, recruiting interneurons and increasing the phase lag between the stimulus and the network; similar responses might be observed in low-acetylcholine states *in vivo*. Importantly, both network states are well-calibrated to *in vivo* data, suggesting that these state-dependent effects are indeed biologically relevant. Surprisingly, however, recurrent connectivity does not amplify the entrainment of pyramidal cells, rather acting as a baseline offset to network activity. Together, these results shed light on the mechanisms of tACS at the circuit level, and constrain how stimulation should be coupled to reduced models of cortical tissue: the effective sensitivity of a circuit to a weak field, and the phase of its response, depend critically on stimulation frequency and on network state.

## 2 Results

### 2.1 tACS entrains network activity at realistic field strengths

Simulations were performed using a single cortical column (approximately 30,000 neurons) of the Blue Brain Project’s rat somatosensory model (Fig. 1a; see Section 4.1 for details) [27, 28]. To assess truncation effects stemming from the limited spatial extent of the single column, we compared the single-column response with that of a larger cortical subvolume comprising seven contiguous columns (approximately 210,000 neurons) under 10 Hz stimulation at various amplitudes. Entrainment magnitudes were similar in the two circuits, although their phase distributions differed slightly at the highest studied field magnitude of 4 V m*^−^*^1^ (Supplementary Section S1 and Supplementary Fig. S.1). As the network response did not differ substantially between the two different circuit sizes, we used the single-column circuit for the remaining simulations to limit computational cost.

Endogenous network synchrony is thought to play a role in shaping the response to stimulation [12, 13, 17, 33]. To investigate this hypothesis, we simulated the network in two characteristic modes: a “REM-like” state, in which the firing activity of excitatory and inhibitory populations is relatively desynchronized, and a “NREM-like” state, with increased correlation between the spike-time histograms of excitatory and inhibitory populations. We use the terms “REM-like” and “NREM-like” as rough phenomenological descriptions of network activity—not to indicate that the specific neurobiology generating the different states is simulated (for further details see Section 3.3.1). The network state was controlled by injecting Ornstein-Uhlenbeck conductance noise into the soma of each neuron in the circuit, with a fixed amplitude per neural population. These amplitudes were determined by an algorithm [28] that optimizes model spike rates against experimental data from an anesthetized animal; the two states differ in extracellular calcium concentration (which is associated with synaptic efficacy [34]), in the input-noise ratio R*_OU_* , and in the firing-rate calibration target (see Section 4.1). In the absence of electrical stimulation, the NREM-like network produces a strong oscillation at 2.5 Hz, in contrast to the intrinsic activity of the REM-like network, which has much smaller peaks of oscillatory power at 2.5 Hz, 7.5 Hz, 10 Hz, and 15 Hz (Supplementary Fig. S.2). The increased delta-frequency power in the high-correlation state, together with the calibration of injected noise to *in vivo* activity under urethane anesthesia, which is known to preserve naturalistic sleep states [35], motivates the use of the term “NREM-like” for the high correlation condition, and “REM-like” for the low-correlation condition.

We analyzed the response of the network to spatially uniform extracellular electrical stimulation oriented along the depth axis of the column, at frequencies of 7 Hz, 10 Hz and 15 Hz, (corresponding to the theta, alpha, and beta ranges, respectively; 15 Hz stimulation was performed only in the REM-like state) and at electric field intensities ranging from 0.25 V m*^−^*^1^ to 4 V m*^−^*^1^. Stimulus amplitudes in the lower portion of this range (<1 V m*^−^*^1^) are typical of human tACS [36, 37], while the upper portion extends into the range of field strengths typical of animal experiments (where peak fields can even be much higher than those studied here) [38, 39] and computational studies [17]. The applied fields were in the subthreshold regime, well below the lowest reported threshold for direct action potential initiation (approximately 12 V m*^−^*^1^) [30]. In response to 1 second of stimulation, mean firing rates changed little relative to baseline for stimulus amplitudes up to 2 V m*^−^*^1^. At 4 V m*^−^*^1^, firing rates increased in both network states. The degree to which firing rate was elevated under 4 V m*^−^*^1^ stimulation varied across stimulus frequencies, ranging from 9.5% to 16% in the REM-like state; in the NREM-like state, firing rates increased by 43% for 10 Hz stimulation and 87% for 7 Hz stimulation (Supplementary Fig. S.3). To measure the effects of stimulation on endogenous oscillations, we apply 2 seconds of 10 Hz stimulation at 1 V m*^−^*^1^ to the REM-like-state network (note, however, that throughout the rest of this paper stimulation is applied for only 1 s). While endogenous oscillatory power is weak in this condition, stimulation nevertheless substantially reduced oscillatory power at off-target frequencies, while power at the target frequency was substantially enhanced (Supplementary Fig. S.4a). At most frequencies, power is reduced in the second half of the 2 s stimulation window compared to the first half (Supplementary Fig. S.4b), which may reflect vesicular depletion during stimulation, a possibility not directly tested here.

We quantified the entrainment of network activity to the stimulus with the phase-locking value (PLV), a measure of the consistency of the phase difference between the stimulus waveform and spiking activity in the network (see Fig. 1c-h and Section 4.5.1). Network PLV varied with stimulus amplitude, frequency, and network state (Fig. 2a). To account for the finite-spike-count bias of the squared PLV, we also calculated pairwise phase consistency (PPC; see Section 4.5.1). Across the frequency-state combinations analyzed here, PPC increased with stimulus amplitude, with the slope of the increase varying with frequency; at a given amplitude, stimulation with lower frequencies tended to produce greater PPC than stimulation at higher frequencies (Fig. 2b). Because PPC approximates the squared PLV, the supralinear increase of PPC at amplitudes above roughly 1 V m*^−^*^1^ corresponds to an approximately proportional increase of PLV with amplitude (Fig. 2a,c). However, for lower amplitudes, nonlinear increases in PLV were observed for 10 Hz stimuli, particularly in the REM-like state (Fig. 2c). For stimulus frequencies of 7 Hz and 10 Hz, significant differences from baseline in preferred-phase distributions were not observed below a stimulus amplitude of 1 V m*^−^*^1^ in either the REM-like or NREM-like state, while for 15 Hz stimulation in the REM-like state, a significant difference was detected only at 4 V m*^−^*^1^ (Fig. 2e). Significant differences in preferred-phase distributions between 7 Hz and 10 Hz stimulation in both network states, and between 7 Hz and 15 Hz in the REM-like state, were observed for a stimulus amplitude of 4 V m*^−^*^1^ (Fig. 2e).

**Figure 2:**
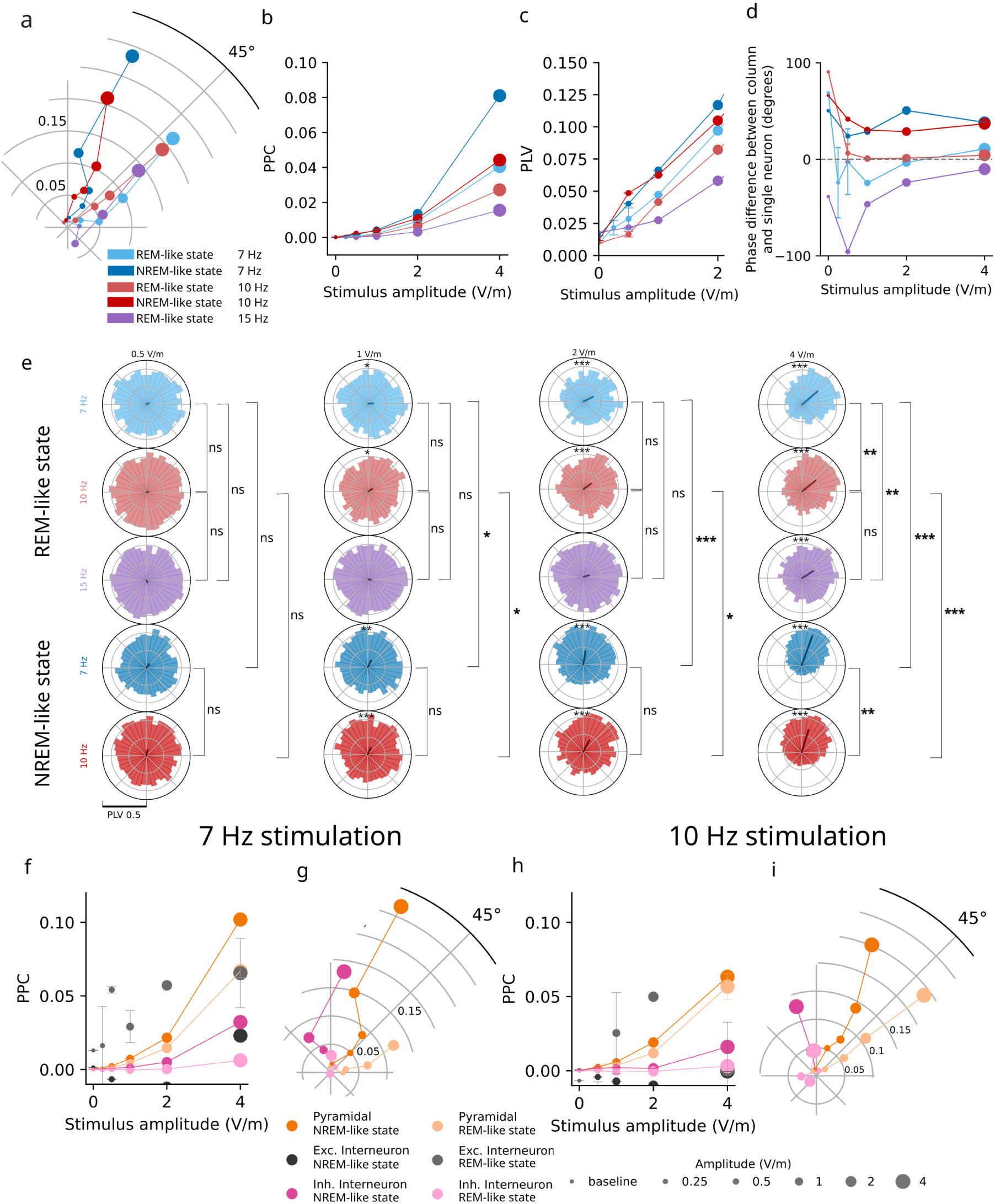
Network entrainment in response to extracellular stimulation. a: Phase-locking value (relative to the stimulus waveform) of the firing activity of the network, as a function of stimulus amplitude (represented by marker size), frequency (represented by color), and network state (represented by lighter and darker shades). Radius corresponds to degree of entrainment; angle corresponds to phase lag, with larger angles indicating greater delay. b-c: Pairwise phase consistency (b) and PLV (c) increase with stimulus amplitude across frequencies and network states. Error bars represent standard deviation over multiple simulations with the same stimulus and network state, but varying random seeds. d: In the REM-like state, but not the NREM-like state, as stimulus amplitude increases, the network phase lag approaches the phase lag of the somatic polarization of a single layer 5 pyramidal cell exposed to an extracellular field. e: Histograms of cells’ phase preferences, for each combination of stimulus frequency, amplitude, and network state. Asterisks on phase preference histograms indicate significant differences from the unstimulated network in the same state. Brackets indicate pairwise comparisons of preferred-phase distributions between frequencies within a state and between states at the same frequency (Moore’s paired test; No asterisk or ns: p > 0.05, *: p < 0.05, **: p < 0.005, ***: p < 0.0005). f-g: PPC (f) and PLV (g) for different cell types (indicated by color) and different network states (indicated by shade) in response to 7 Hz stimulation. Error bars represent standard deviation over multiple simulations with the same stimulus and network state, but varying random seeds. h-i: As in panels f-g, but for 10 Hz stimulation.

In addition, we stratified the stimulation response by cellular identity, dividing the population into pyramidal cells, inhibitory interneurons and excitatory interneurons, and assessing entrainment accordingly. As stimulus amplitude increased, a more pronounced increase in the PPC of pyramidal cells, compared to other cell types, was evident for all network conditions (Fig. 2f,h), though state-dependent differences in the response of inhibitory interneurons were also observed (see Section 2.2). While excitatory interneurons sometimes entrained strongly to the stimulus (Fig. 2f,h), this response was highly variable, non-systematic (see Supplementary Fig. S.5, S.6), and difficult to interpret due to the small total number of spikes. In contrast, the observed trends in network-level PPC are closely paralleled by the PPC of pyramidal cells.

### 2.2 Entrainment of network activity depends on network state

The NREM-like state was generated using a higher extracellular calcium concentration and a lower input-noise ratio R*_OU_* than the REM-like state (see Section 4.1); it therefore has greater synaptic efficacy and is driven more by internal connectivity and less by noise. At both 7 Hz and 10 Hz, the NREM-like network exhibited stronger entrainment to the stimulus than the REM-like network (Fig. 2a-b; 15 Hz stimulation was not examined in the NREM-like state). Statistically significant differences in preferred-phase distributions between the REM-like and NREM-like states were observed at all stimulus amplitudes 1 V m*^−^*^1^ (Fig. 2e). Compared to the REM-like network, stronger phase locking of inhibitory interneurons was observed in the NREM-like network (Fig. 2e-h), with a significant phase preference beginning at 2 V m*^−^*^1^ for 7 Hz stimulation (p < 0.0005) and at 4 V m*^−^*^1^ for 10 Hz (p < 10*^−^*^5^) (Supplementary Fig. S.6c).

While the phase of the network activity at the stimulus frequency was effectively random for low-amplitude stimulation (Fig. 2a,c), for higher-amplitude stimulation in the REM-like state, the phase synchronized to 40*^◦^* for all frequencies (Fig. 2a). As previously described, prior modeling work has established a clear relationship between tACS and somatic polarization in individual cells. We therefore compared the phase of the network response to tACS to the phase response of somatic polarization in an isolated layer 5 pyramidal cell (see Section 4.3). It should be noted that the two metrics are not directly comparable; the network phase lag reflects the delay between the stimulus and *spiking activity*, and partially reflects synaptic processes, while somatic polarization is driven entirely by the electronic structure of the cell. However, as somatic polarization does directly influence the probability of action potential generation, the comparison between the two metrics may be informative. We found that the network phase delay in the REM-like state approached the somatic polarization phase lag of the isolated cell as stimulus amplitude increased (Fig. 2d), suggesting a shift from intrinsic network oscillations toward direct stimulus-driven entrainment. Interestingly, and in contrast to the REM-like circuit, the phase lag of the network activity in the NREM-like state did not converge to the phase lag of an isolated single cell (Fig. 2a,d), suggesting that strong stimulation in the presence of enhanced synaptic efficacy recruits local polysynaptic pathways. Moreover, the phase lag of the NREM-like network was substantially larger than that of the REM-like network, indicating a more delayed response to stimulation. This further supports the hypothesis that poly-synaptic pathways may mediate the response of the NREM-like network, as a longer synaptic pathway would naturally delay the response of the network.

### 2.3 Recurrent feedback shapes the preferred phase of the network response in the NREM-like state

To assess the contribution of stimulation-dependent recurrent feedback to the circuit response to tACS, we compared the connected network with a replay condition, referred to as the “disconnected network”, in which neurons receive only noise input and the synaptic input recorded from an unstimulated connected simulation (i.e., their synaptic input from the other neurons in the simulation is not affected by the electric exposure), such that stimulation-induced changes in spiking cannot feed back through the circuit (Fig. 3a; see Section 4.4 for details). During 10 Hz stimulation at 2 V m*^−^*^1^, no substantial differences were observed in entrainment amplitude between the connected and disconnected conditions, in either the REM-like or NREM-like network states (Fig. 3b). These results indicate that recurrent connectivity is not required to produce neural entrainment under realistic exposure conditions and that network connectivity does not amplify the impact of electric fields on neural entrainment. Similarly, no substantial differences in entrainment amplitude were observed for pyramidal cells specifically (Fig. 3c); while differences did appear for inhibitory interneurons, differences in inhibitory interneuron phase preference distributions rose to statistical significance only for the NREM-like state (Fig. 3d).

**Figure 3:**
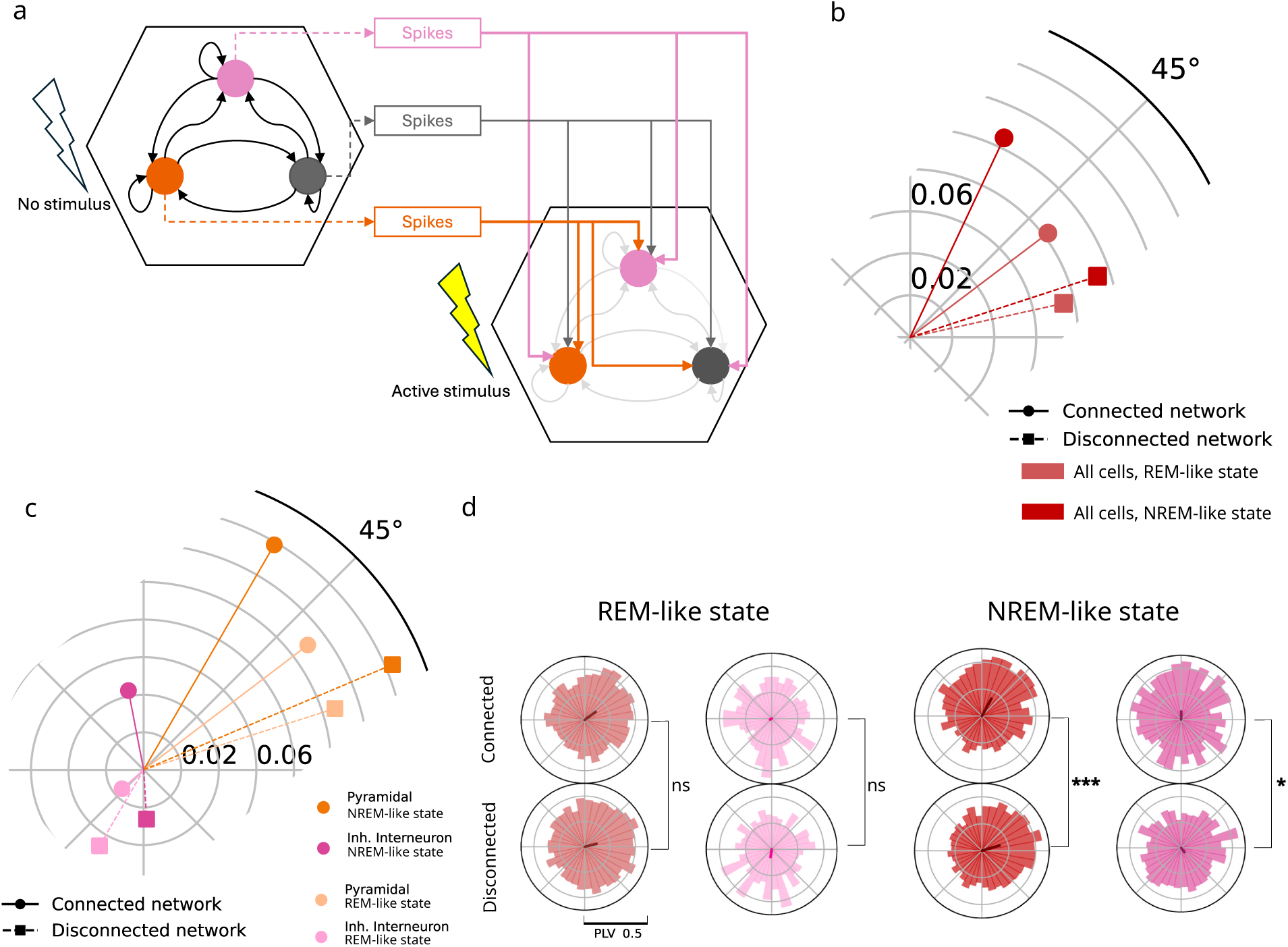
Recurrent feedback shapes the phase of the network response. a: Schematic of the spike replay protocol, adapted from [40]. Circles represent neural populations (for clarity, only three are sketched). Black arrows represent active synaptic connections within the circuit; gray arrows represent inactive connections within the circuit. Dashed colored arrows represent the recording of spikes from each population in the connected network, while solid colored arrows represent the artificial activation of efferent synapses from these populations in the disconnected circuit. Lightning bolt symbols represent the absence (unfilled) or presence (filled) of extracellular stimulation. b: Resultant vectors of all network spikes during 10 Hz stimulation at 2 V m*^−^*^1^. Stimulation-dependent recurrent feedback has little effect on the PLV magnitude in either state; it shifts the preferred phase of the network response modestly in the REM-like circuit and substantially in the NREM-like circuit. c: As in b, for pyramidal cells and inhibitory interneurons. The preferred phase of both populations is significantly affected by recurrent feedback in the NREM-like, but not the REM-like, network state, with little change in the PLV magnitude of pyramidal cells. d: Significant differences in preferred-phase distributions are observed for all cells in the network, and specifically for inhibitory interneurons, between the connected and disconnected conditions in the NREM-like state, but not in the REM-like state.

In the REM-like state, the network preferred phase shifted by approximately 25*^◦^* between the connected and disconnected conditions (Fig. 3b). This shift was not statistically significant at the level of individual neurons’ preferred phases (Fig. 3d); no large differences between the two conditions were observed in any neural subpopulation in this state (Fig. 3c). In contrast, the phase response of the NREM-like circuit differed significantly between the connected and disconnected conditions (Fig. 3b,d): overall phase was shifted by 47*^◦^*, while pyramidal cell phase across all layers was shifted by 37*^◦^*. Statistically-significant differences in phase preferences were observed both for the network as a whole, and for inhibitory interneurons specifically (Fig. 3d).

Interestingly, we observed that in both the REM-like and NREM-like states, pyramidal cells consistently have a smaller phase lag than inhibitory interneurons. Moreover, we found that disconnecting the network reduces the phase lag of pyramidal cells, with variable effects on the phase lag of inhibitory interneurons—this trend also held for most layer-resolved populations of pyramidal and inhibitory neurons (Fig. S.7).

## 3 Discussion

In this study, we analyzed the response of a detailed model of a rat somatosensory cortical column [28] to subthreshold tACS stimulation, applied as a uniform field along the main axis of the column. To our knowledge, this is the first demonstration, in such a detailed model of a cortical circuit, that tACS applied at realistic amplitudes entrains neural oscillations. Nonlinear, frequency- and brain-state-dependent network responses to tACS arise from our model as emergent phenomena, even though the model was built from the bottom up without any manual tuning. The simulations show that stimulus locking depends on field amplitude, frequency, and simulated network state, while spike-replay comparisons indicate that stimulation-dependent recurrent interactions shape the preferred phase of the response, particularly in the NREM-like state.

### 3.1 Network response is more complex than predicted by single-neuron models

#### 3.1.1 Network entrainment depends non-trivially on stimulus amplitude and frequency

Intensity-dependent effects of tACS have been reported in simplified network models [17], in a small morpholog- ically detailed microcircuit [25], in single-cell models [6, 7], and in single-unit recordings from awake nonhuman primates [10, 11, 33], but had not yet been examined in a model of this size and anatomical detail. Zhao et al. [17] constructed a computational network model of two-compartment pyramidal neurons and inhibitory interneurons, but did not capture the morphological and connectomic details and *in vivo*-like endogenous ac- tivity which shape responses in our circuit. Park et al. [25] modeled entrainment with morphologically detailed neurons, but using only one representative neuron per cortical layer. In contrast, with 30,000 morphologi- cally detailed neurons, our model captures a significantly greater share of the cortical column’s full biophysical complexity.

Single-cell models have established an approximately linear relationship between stimulus amplitude and entrainment [6, 7], though how this translates to a network context is unclear. In our network model, we observe monotonic increases in entrainment with electric field intensity; PPC, which approximates the square of PLV, increased supralinearly with amplitude, whereas PLV increased approximately in proportion to field strength for high stimulus amplitudes (Fig. 2a-b). However, the relationship between PLV and stimulus amplitude appeared more variable for low field strengths (Fig. 2c). Prior work has suggested that low-amplitude stimulation at, or slightly offset from, endogenous frequencies can disrupt phase locking, while higher-amplitude stimulation reinstates it with a different phase preference [11, 17, 25]. At low stimulus amplitudes, the preferred-phase comparisons did not detect significant changes relative to baseline (Fig. 2e); the absence of a detectable disruption may reflect the fact that, in comparison to previous computational work [17, 25], our model was built from the bottom up, rather than tuned to produce a specific endogenous oscillatory frequency. This may lead to lower baseline PLVs than those observed in prior studies. Experimental evidence suggests that stimulation-induced reductions in phase locking are strongest in neurons with the highest baseline phase locking [11]; the relatively weak baseline locking in our model may therefore limit our ability to detect such reductions. We further observed a frequency-dependence in the REM-like network response to stimulation, where the network entrained more efficiently to frequencies of 7 Hz and 10 Hz than to 15 Hz (Fig. 2a,b,d). The power spectrum of the spontaneous network activity revealed that, though the REM-like network did not have any particularly strong oscillation frequency, the network did exhibit relatively small peaks of oscillatory power at 2.5 Hz, 7.5 Hz, 10 Hz and 15 Hz (Supplementary Fig. S.2b). While power exactly at the stimulus frequencies was comparable in all cases, more power was present in the regions around 7 Hz and 10 Hz than in the region around 15 Hz. Prior work has shown that network responses to tACS follow an Arnold tongue, such that stimulating at, or close to, the endogenous frequency of the network induces the strongest entrainment [12, 13, 17, 33]. The weaker entrainment at 15 Hz is compatible with such a frequency-selective response.

#### 3.1.2 Impact of network state on entrainment is not predictable from baseline activity

We next simulated the network in a NREM-like state, in which the circuit exhibits strong oscillatory activity and increased synaptic efficacy. The NREM-like network state had a stronger response than the REM-like state at the frequencies examined in both states (7 Hz and 10 Hz) for amplitudes 1 V m*^−^*^1^ (Fig. 2a-d). The difference in locking strength was not predicted by baseline power at the stimulation frequencies alone—power at 7 Hz and 10 Hz was similar in the two states (Supplementary Fig. S.2c). However, stronger locking is consistent with the increased synaptic efficacy in the NREM-like state (though, as the difference in entrainment between the two states is not abolished in the disconnected network condition (Fig. 3b), mechanisms other than recurrent feedback must also play a role; see also Section 3.2.1). In contrast with our observation that the network exhibits more entrainment in the NREM-like state compared to the REM-like state, previous modeling work by Taghavi et al. has suggested that, in comparison to networks with lower endogenous coherence, networks with high endogenous coherence exhibit weaker entrainment for stimulus frequencies distant from the endogenous oscillation frequency [41]. While Taghavi et al. generated endogenous oscillatory activity using Poisson noise input [41], the endogenous oscillations of our network depended on a biologically motivated choice of meta- parameters, including extracellular [Ca^2+^]. This suggests that whether circuits display a strong preference for stimulation at the endogenous frequency may depend on the mechanisms by which different network states are instantiated. We therefore propose that a characterization of network state that extends beyond oscillatory power at stimulus frequencies is necessary in order to predict network responses to tACS.

We also observe differences in the network phase response between the REM-like and NREM-like states. In the former, we found that, with increasing stimulus amplitude and for all stimulus frequencies, the phase lag of the network converged to a common value similar to the phase lag of an isolated pyramidal neuron (Fig. 2d). In contrast, the phase response in the NREM-like state was substantially larger than that of an isolated neuron (Fig. 2d). One interpretation is that the initial, polarization-facilitated spiking increase recruits additional, delayed activity through recurrent polysynaptic pathways, whose efficacy is higher in the NREM- like state. Within each cycle, this recruitment continues either until the network settles at a steady level of activity, at which the recurrent input generated by the population’s own firing is just sufficient to sustain it, or until the declining phase of the alternating field reduces the polarization and, consequently, the spiking. As a result, more time is needed for the network to build to its peak response. This interpretation also aligns with the layer-dependence of the phase delay apparent in Figure S.7 that vanishes in the disconnected network: layer 4, and to a lesser degree layer 5/6, function as the primary input layers for feed-forward information processing (n.b., extra-columnar input was modeled here as noisy conductance). Subsequently, layer 4 projects to layer 2/3 layer 5 layer 6, introducing asymmetry in the distribution of synaptic connections. When an external field is applied, this asymmetry is reflected in the sequence of polysynaptic pathway activations and thus the phase of maximal layer entrainment, resulting in a similar response sequence. Characterizing this state- and layer-specific phase lag may enable closed-loop targeting of the ongoing oscillatory activity of specific neural populations, particularly during stimulation in NREM sleep, where the phase delay is greater. The increased phase delay of the NREM-like network is not predictable from baseline spectral power at the stimulation frequency (a typical metric to characterize network state in the context of tACS), highlighting the need for a more comprehensive description of network state; relatively higher PSD peak and baseline firings rates (Supplementary Figs. S.2 and S.3, respectively) might be useful markers of the connectivity-mediated network coherence underpinning these state-dependent phase differences.

### 3.2 Towards a mechanistic understanding of tACS

Our results suggest that the network response to stimulation is initiated, and primarily driven by, direct en- trainment of pyramidal neurons to the extracellular field. Depending on network state, synaptic input from pyramidal neurons to inhibitory interneurons may entrain the latter. Similarly, synchronized recurrent input to pyramidal cells may, depending on network state, shift their spike timings, without a substantial impact on the magnitude of entrainment to the stimulus.

Several lines of evidence support this interpretation. In all network states, the response of pyramidal cells to the stimulus is stronger, and has a smaller phase lag, than the response of inhibitory interneurons (Fig. 2f-i). Critically, disconnecting recurrent input does not have a substantial impact on the magnitude of pyramidal cell entrainment (Fig. 3c), confirming that they are directly entrained by the extracellular field. That pyramidal cells are directly entrained is largely accepted, though some recent studies have found greater entrainment in inhibitory cells than pyramidal cells [13].

Unlike pyramidal cells, substantial differences in the magnitude of inhibitory entrainment are observed between the connected and disconnected conditions (Fig. 3c). While these differences are difficult to interpret due to the opposite direction of change in the REM-like versus NREM-like states, we suggest that inhibitory interneurons are entrained by synaptic input in the NREM-like but not the REM-like state. This hypothesis is consistent with the observation that inhibitory entrainment increases more steeply with stimulus amplitude in the NREM-like state than in the REM-like state (Fig. 2)f-i—this difference is attributable to increased synaptic efficacy. Moreover, inhibitory phase lags behind that of pyramidal cells for all stimulus amplitudes, with a relatively consistent offset (Fig. 2g,i), implying synaptic, rather than direct, entrainment. (Note that the inhibitory phase offset differs sharply for 7 Hz stimulation at 4 V m*^−^*^1^; in this condition, firing rates were substantially increased (Supplementary Fig. S.3b), complicating interpretation.) It may be that in the REM-like state, pyramidal input desynchronizes inhibitory interneurons due to weak synaptic efficacy; however, given the phase variability and weak entrainment observed (Fig. 2f-i), it is also possible that reduced entrainment in the disconnected condition is attributable to random variation. Several previous studies [6, 13, 25] have suggested that inhibitory neurons are entrained by synaptic input; our results caveat this observation, finding that such entrainment is network-state dependent.

Unlike previous modeling work from Park et al. which found an important role for synaptic connectivity in the phase response of interneurons only [25], our data suggests that synaptic connectivity plays an important role in determining the network-state-dependent phase response of pyramidal cells as well. As previously described, the phase lag of our NREM-like network was substantially larger than that of the REM-like network. However, this difference is largely abolished in the disconnected state (Fig. 3b). In contrast, recurrent connectivity has a smaller effect on the phase-lag of the REM-like network (Fig. 3b). While the existence of such a shift might suggest that the phase response of the NREM-like network is partially driven by recurrent connectivity, this shift does not rise to the level of statistical significance (Fig. 3d); that the phase response of the NREM-like network more closely approaches the phase of somatic polarization in an isolated neuron in the connected condition than the disconnected condition (Fig. 2d) may be an artifact of the choice of a particular neuron in the isolated condition. Thus, we conclude that the phase response of the network depends on both network state and connectivity. As Park et al.’s circuit comprises only five synaptically connected neuron models, with a single representative cell per cortical layer, and as network activity was tuned to produce a single narrow spectral peak rather than the broadband activity characteristic of *in vivo* cortex [25], it is not surprising that our modeling results diverge, though we did not disambiguate differences due to model size, background activity, and connectivity. To our knowledge, network-state- and connectivity-dependence in the network phase response have not previously been demonstrated in detailed network models.

#### 3.2.1 Cellular and circuit mechanisms underlie network state dependence

We have shown that network state affects both the magnitude of entrainment to stimulus, and the phase response of the network. While our results suggest that recurrent connectivity is responsible for the phase shift in the NREM-like state, it does not establish whether inhibitory input is necessary, or if the effect is due to recurrent excitatory input. Previous computational modeling has suggested that excitatory synaptic input increases phase lag, while inhibitory input reduces it [17], and that excitatory synaptic input enhances entrainment, while inhibitory input weakens it. In our model’s NREM-like state, we observe a slight increase in the magnitude of pyramidal entrainment in the disconnected versus connected condition (Fig. 3c). It may therefore be the case that both excitatory and inhibitory recurrent connectivity drive responses, with excitatory recurrent connectivity shifting the phase, and inhibitory input canceling the effects of excitatory input on entrainment magnitude. However, Zhao et al. study the effect of connectivity on phase response in an isolated pair of neurons with feedforward connectivity only [17]. Thus our model may capture more complex behavior emerging from recurrent, poly-synaptic connectivity, which was not present in prior work.

We found that the difference in entrainment amplitude between network states is somewhat reduced, but not entirely eliminated, in the disconnected condition (Fig. 3b); differences in synaptic efficacy between the two network states therefore only partially account for differences in entrainment. As previously discussed, the entrainment magnitude of pyramidal cells does not differ between the connected and disconnected conditions (Fig. 3c), and the difference in entrainment between the REM-like and NREM-like states is partially explained by the increased synchronization of inhibitory interneurons via synaptic input in the NREM-like state. The remaining entrainment increase in the NREM-like state must therefore be due to increased entrainability at the single-neuron level, which might arise due to increased baseline firing rates (Supplementary Fig. S.3) and reduced variance in input noise (see Section 2.2).

#### 3.2.2 Connectivity acts as an offset, not an amplifier

While it is not surprising that pyramidal cells are directly entrained by the electric field, it is unexpected that network connectivity does not enhance, and may even disrupt, their entrainment (Fig. 3c). While this result is consistent with some prior observations [15], it differs from studies reporting stronger recurrent contributions to entrainment magnitude [12, 13]. Indeed, network-level amplification of neural entrainment has been considered one of the more likely potential mechanisms by which very weak somatic polarization could result in detectable network responses [42]. Rather, in our model, neurons receive sufficient input at baseline such that even a small somatic polarization suffices to induce changes in spike timing. That the degree of entrainment does not differ substantially under the connected and disconnected conditions does not imply that network connectivity plays no role in the response to stimulation; it is precisely the replay of synaptic input from the baseline simulation that brings neurons into a stimulus-sensitive state. The spike replay technique used here also preserves the spatial arrangement of synaptic inputs across the neuronal arbor, which may interact with the electric field in complex ways [18]; the role of the spatial distribution of synapses in the network response should be explored in future work.

It is likely that differences in network state explain the divergence between our findings and previous work. In [12], the network was placed, by design, in a state that was far more synchronous than even our NREM-like state. This extreme synchrony likely enhanced excitatory-excitatory synaptic mechanisms underlying entrainment in the model—to a greater degree than could be expected in most biological network states. Similarly, Huang et al.’s finding that indirectly-recruited inhibitory interneurons are more strongly entrained than pyramidal cells [13] likely depends on brain state. Their experimental results uncovered much stronger phase locking in the posterior parietal cortex—a region that is likely the source of the endogenous alpha oscillations targeted in that work—than in other regions with similar tACS field exposure. It should also be noted that the computational models used by Ali et al. [12] and Huang et al. [13] featured networks of point neurons with limited variability, more prominent intrinsic oscillations, and a less complex connectivity structure than the circuit model employed here. Given previous observations that excitatory input tends to increase PLV and inhibitory input tends to reduce it [17], we hypothesize that our network would produce network amplification of tACS if it were placed in a state with less robust E/I balance. We note, however, that the network states studied in this work are well-calibrated to *in vivo* data; any future exploration of alternative network states should be careful to maintain this connection with *in vivo* biology.

### 3.3 Limitations of the study

Despite its cellular and circuit detail, the model retains several simplifications that may influence its responses to stimulation. The simulated cortical column has a radius of approximately 230 µm , substantially smaller than the tissue volume exposed during tACS. While single- and seven-column circuits showed similar entrainment magnitudes under the tested 10 Hz conditions, with only a slight difference in their phase distributions at 4 V m*^−^*^1^ (Supplementary Fig. S.1), the influence of longer-range connections, from regions sufficiently distant as to experience a non-uniform stimulus field, remains to be established. While modeling such a large volume of tissue is currently infeasible from a computational perspective, future advances may enable the impact of long-range connectivity to be studied in further detail.

In addition to the network size, several aspects of neural physiology were simplified for computational efficiency. In particular, axons are truncated to an axon initial segment of 60 µm and a myelinated segment of 1 mm [34]. Axonal morphology and physiology influence direct activation by extracellular stimulation [30], but their relevance is not restricted to suprathreshold stimulation; experiments in cultured neurons have shown that subthreshold direct-current fields can alter neurotransmitter release through presynaptic polarization [43]. Those experiments used direct-current fields of 5 V m*^−^*^1^ and above, beyond the upper end of the range studied here, and the contribution of this mechanism to tACS responses remains untested. Future work should assess the sensitivity of the model to axonal morphology and physiology.

*In vivo*, cells are affected not only by synaptic interaction and exogenous electric fields, but also by endogenous electric fields generated by neural activity, both in other segments within the cell and by nearby cells in the network (co-called “ephaptic coupling”). Endogenous fields of 2 V m*^−^*^1^ to 4 V m*^−^*^1^ have been measured during *in vivo* slow oscillations, and applied fields of a similar magnitude (comparable to the upper end of the range studied here) can guide network activity [44]. Our model does not simulate ephaptic coupling, as computing the effect of transmembrane currents on every other compartment in the network would be computationally infeasible. Since ephaptic coupling synchronizes activity between neurons, we hypothesize that its effects on the network response to stimulation will be similar to that of the NREM-like network state. However, as its effects are likely sensitive to the spatial and temporal structure of the endogenous fields, its impact on entrainability in the network states studied here remains unresolved. Future work, computing a mesoscale transfer function between extracellular stimulus and neural response, may be able to account for the effects of ephaptic coupling *post hoc* by applying a similar transformation as that generated by the exogenous field.

In our simulations, we assume that the electric field induced by tACS is spatially uniform across the depth of the cortical column. While historically, the spatial gradient of the electric field has been believed to be locally negligible at the scale of a single cortical column [45], recent work has suggested that tissue conductivities, and consequently electric field strengths, may vary across cortical layers [46]. Variation in electric field strength with depth may have unpredictable consequences for neural responses, particularly for pyramidal cells with dendrites spanning multiple layers of the cortical column. While it is not feasible to explicitly model the conductivity variation caused by individual neurites in the circuit, it may be possible to parameterize the variation in conductivity over the depth of the cortical column at a coarser scale, and thereby account for variation in electric field strengths.

In the present work, we explored only a small portion of the tACS parameter space, importantly, not including amplitudes 0.3 V m*^−^*^1^, which have been reported to disrupt ongoing oscillations in simplified models [17, 25], or frequencies of 30 Hz and above, at which inhibitory interneurons have been shown to entrain more strongly *in vitro* [5]. We therefore expect a more complex network response to stimulation outside of the parameter range studied here. Nevertheless, tACS at relatively low frequencies and high amplitudes is biologically interesting, as many neural oscillations of clinical relevance lie in this frequency range, and *in vivo* tACS can reach the field strengths considered here, particularly, but not exclusively, in animal models. Additionally, stimulation was applied only parallel to the main somato-dendritic axis of pyramidal cells, an orientation that generally produces strong somatic polarization [18, 21, 30]. Other orientations can affect different neuronal compartments and signal propagation [18]; their effects on the network responses reported here remain to be investigated.

A more complete mechanistic understanding of the effects of tACS is limited by the number and type of *in silico* experiments we were able to perform. While we propose circuit-level mechanisms for the neural entrainment observed in this study (Section 3.2), we cannot yet elucidate the specific neural pathways involved. Selective pathway perturbations combined with spike replay (see Tharayil et al. [40]) could help distinguish these contributions. Similarly, while we were able to establish differential responses to stimulation in two network states and suggest mechanistic hypotheses for these differences (Section 3.2.1), we cannot unambiguously confirm these hypotheses, as several parameters were co-varied across states. Future work with a more fine-grained parameter sweep could shed light on these mechanisms.

#### 3.3.1 Use of the terms “REM-like” and “NREM-like”

We refer to the two network states studied in this work, with, respectively, low and high correlation between excitatory and inhibitory activity, as “REM-like” and “NREM-like”. These descriptions are rough, phenomeno- logical descriptions; the model does not replicate the neurobiology of sleep. Nevertheless, we argue that the relationship between the network states in our model and sleep in not merely superficial; the parameters control- ling network state in the model partially recapitulate biological features of different sleep states, so the behavior of the model may provide some insights into *in vivo* behavior in different sleep stages.

While the parameter combinations ([Ca^2+^], P*_F R_* and R*_OU_* ) in both of the simulated network states were selected based on comparisons to *in vivo* activity under urethane anesthesia rather than naturalistic sleep [28], urethane is known to produce oscillations between weakly-correlated, REM-like neural activity and strongly- correlated, NREM-like neural activity [35]. *In vivo*, this transition has been shown to be mediated by an increase in cholinergic tone in the REM-like state [35]. In contrast, our model represents the REM-like state with an increased P*_F R_* and R*_OU_* , and decreased [Ca^2+^], compared to the NREM-like state. These changes parallel some of the alterations induced by cholinergic tone in the cortex: intracortical synaptic efficacy is suppressed by both cholinergic activation of muscarinic receptors *in vivo* [47] and by decreasing [Ca^2+^] in our model [28]; cholinergic activation of nicotinic receptors increases thalamic input to cortex *in vivo* [47, 48], which is paralleled by increased P*_F R_* and R*_OU_* in our model.

### 3.4 Summary and Outlook

Our simulations reproduce several phenomena reported across experimental preparations: intensity- and frequency- dependent entrainment [9, 10, 13, 33], a dominant role of pyramidal cells at low stimulation frequencies [5, 15, 21], and a strong dependence on ongoing network state [11, 49]—all within a single, anatomically and physio- logically constrained model. Because the model allowed us to manipulate the mechanisms underlying different network states and to disconnect the network while replaying its activity, it yielded mechanistic insights that would be difficult to obtain experimentally. For weak stimuli, entrainment depends on stimulus amplitude in a more complex manner than would be predicted by single-cell polarization. We found that network state impacts both the amplitude and the phase of the circuit response to stimulation, and that these impacts can- not be predicted from endogenous activity at the stimulation frequency. We found that recurrent connectivity shapes the phase of the overall response in a state-dependent manner, but that, in the network states studied, recurrent connectivity does not amplify neural responses to stimulation. Together, these results suggest that the coupling between extracellular stimulation and the network response is more complex than the traditional “λ⃗ E⃗” approach to modeling mesoscale stimulation responses would imply. Moreover, the present work lays the foundation for a mechanistic understanding of tACS at the circuit level, and highlights the importance of network state in the response to extracellular stimulation. It suggests that mechanistic evidence developed from *in vivo* studies should be interpreted in the light of the ongoing brain state, and that extension of hypotheses developed in one state should only be extended to others with caution.

Future work should aim towards further developing the mechanistic understanding of the interaction between applied extracellular stimuli and neural responses by using selective perturbations to identify the synaptic pathways implicated, particularly in the NREM-like state. A better understanding of the relationship between ongoing network activity and the preferred phase of the neural response to stimulation may help increase the efficacy of transcranial electrical stimulation during sleep, which, despite much promise, has thus far produced variable results [50, 51]. Beyond developing mechanistic understanding, a long-term goal of this work is to infer an effective mesoscale coupling between extracellular stimulation and the network response. Our results suggest that, rather than the linear functions of stimulus amplitude typically used, this coupling may be better expressed in the form of a complex function of stimulus amplitude, frequency and network state. Future work, using biophysically-detailed simulations of the type described in this paper, together with new optimization tools [24], could help to parameterize such coupling functions. Moreover, the work described here can easily be extended to other transcranial electrical stimulation modalities, such as transcranial direct current stimulation and, most interestingly, temporal interference stimulation [52, 53], whose mechanisms are much less understood than those of traditional transcranial electrical stimulation.

## 4 Methods

### 4.1 Network model

We simulate a seven-column subvolume and a single-column subvolume of the BBP rat somatosensory cortex model [27], which represents the two-week-old rat. The seven-column subvolume has a radius of 700 µm and a depth of 2000 µm, while the single-column subvolume has a radius of 230 µm; both subvolumes contain 60 morphological neuron types [27]. The seven-column subvolume consists of 210,000 multicompartmental neuron models, while the single-column subvolume contains 30,000 multicompartmental neurons, where each compartment represents a small discretized segment of a neurite. While all somatic and dendritic compartments are modeled in detail, axons are truncated to the axon initial segment, represented by 10 segments totaling 60 µm in length, and a single myelinated section of 1 mm length. Local connectivity between neurons is defined by an algorithm that identifies potential synapses at appositions between neurons, and prunes these synapses according to biological rules [54]. All simulations are conducted in the single-column subvolume, unless otherwise noted. Ornstein-Uhlenbeck conductance noise is injected in order to simulate missing synaptic inputs. Meta- parameters of the model are varied in order to allow for the simulation of different network states; the input noise is recalibrated for each set of meta-parameters [28]. One such meta-parameter is the ratio of the standard deviation to the mean of the input conductance noise, R*_OU_* = σ*_OU_* /µ*_OU_* , which is fixed across all populations for a given network state. The noise input is optimized to produce *in silico* firing rates equal to a fraction, denoted P*_F R_*, of recorded *in vivo* firing rates [28], to correct for sampling bias introduced by the fact that extracellular recordings preferentially sample more active neurons, thus biasing estimates of population firing rates upward [55, 56, 57].. P*_F R_* is also treated as a meta-parameter. A final meta-parameter is the extracellular calcium concentration (denoted [Ca^2+^]), which ranges from 1.05 mmol L*^−^*^1^ to 1.1 mmol L*^−^*^1^; increasing [Ca^2+^] increases the efficacy of synaptic transmission [28].

In this work, we simulate two different network states, which we denote the “REM-like” and “NREM-like” states, respectively. In the REM-like state, firing activity of excitatory and inhibitory populations is relatively desynchronized, whereas the “NREM-like” state has greater correlation between excitatory and inhibitory firing, resulting in more prominent low-amplitude oscillatory activity at baseline (Fig. S.2). Network states are set by their respective values of [Ca^2+^], P*_F R_* and R*_OU_* ; values for each state are given in Table 1.

**Table 1:**
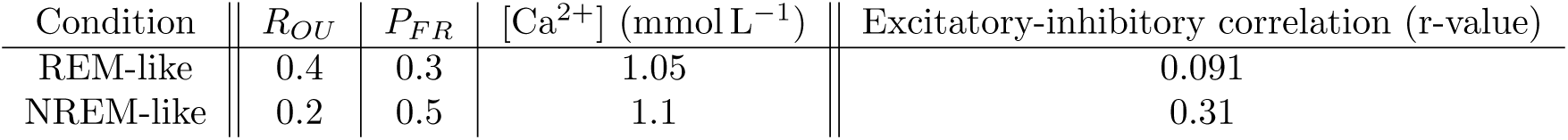
Metaparameter combinations associated with the two network states simulated in this work. Correla- tion coefficients reproduced from [28].

### 4.2 Simulated electrical stimulation

tACS is modeled as a spatially uniform electric field, oriented parallel to the main axis of the cortical column, whose amplitude varies sinusoidally in time, E(t) = A sin (2πf*_s_*(t t_on_)) for t t_on_, with onset at t_on_ = 1500 ms and no amplitude ramp. Stimulation is applied for 1 s, except in the experiment comparing the PSD of network activity during stimulation to baseline, where stimulation is applied for 2 s (Supplementary Fig. S.4). The electric field is coupled to the neuronal models by means of the e extracellular mechanism in the NEURON simulator: at each neural segment, an extracellular potential is calculated by integrating the electric field over the distance between the soma and the segment (note, however, that the choice of the soma as the reference point is arbitrary, as only the difference in the extracellular potential between compartments, and not the absolute value of the potential, affects neural behavior). This calculation is performed at run-time by the Neurodamus simulation control tool [58].

### 4.3 Simulation of isolated cells

We simulate the somatic polarization of an isolated layer 5 pyramidal cell, receiving neither synaptic nor noise input, in response to a spatially-uniform extracellular electric field oriented along the somatodendritic axis. The neuron is a modification by Aberra et al. [30] of a layer 5 pyramidal cell drawn from the circuit, which restores the axonal arbor and increases the size of the neuron; the modifications of [30] have only a minor impact on the phase response of the individual neuron. The particular model used was selected arbitrarily. Somatic polarization is calculated for fields at 7 Hz, 10 Hz, and 15 Hz, and the phase difference between the stimulus waveform and the somatic response is computed. Simulations are performed with a field strength of 1 V m*^−^*^1^; the phase response does not differ substantially with stimulus amplitude. Simulations are performed in Sim4Life v9.0 (ZMT Zurich MedTech AG), using the Electro-Ohmic Quasi-Static solver to calculate the electric field, and the T-Neuro solver to calculate the somatic polarization of a neuron exposed to the field.

### 4.4 Disconnected network simulations

To assess the contribution of stimulation-dependent recurrent feedback to the stimulation response, we used a spike-replay protocol (see Fig. 3). We first recorded all spikes from a connected network simulation without extracellular stimulation. During the subsequent stimulated simulation, synaptic events were triggered by these prerecorded presynaptic spikes rather than by spikes generated online, without altering the transmembrane potential of the presynaptic cell. Neurons therefore continued to receive synaptic drive derived from baseline network activity, while stimulation-induced changes in their spiking could no longer feed back through the circuit. We refer to this replay condition as the “disconnected network”. Stochastic processes, such as sponta- neous vesicle release and conductance noise input , are held identical between fully-connected and disconnected simulation pairs to ensure that cell-cell connectivity is the sole independent variable.

### 4.5 Data analysis

#### 4.5.1 Quantification of entrainment

Neural entrainment was quantified using the phase-locking value (PLV) [59] and pairwise phase consistency (PPC) [32]. Both metrics were calculated directly from neuronal spike times.

For each spike occurring at time t*_j_*, the corresponding phase of the applied stimulation waveform was determined as

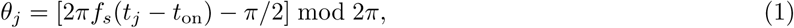

where f*_s_* is the stimulation frequency, and t_on_ is the stimulation onset time.Here, θ*_j_* indicates the peak-to-peak phase difference between the stimulation waveform and the spiking activity. For a set of N spike phases, the mean resultant vector is

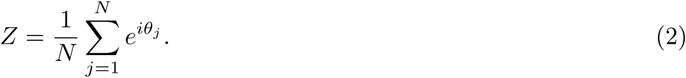

The phase-locking value is PLV = Z , and the preferred phase is θ^^^ = arg(Z), which specifies the lag of the response with respect to the stimulus waveform. PLV ranges from 0 to 1, with larger values indicating that spikes occurred more consistently at a particular phase of the stimulation cycle. Both PLV and θ^^^ can be calculated for the entire network, a neural subpopulation, or an individual neuron. Phase distributions for each *in silico* experiment were calculated by computing the preferred phase θ^^^ for each neuron as described above and binning the resulting values; we use a bin size of 10 degrees.

For small spike counts, as occurs for excitatory interneurons, PLV has an upward finite-sample bias whose magnitude depends on the spike count and on the*_√_*<u>un</u>derlying phase distribution [31]; for uniformly distributed phases, its expected value is approximately 0.89/ N (0.13 for N = 50 and 0.01 for N = 6,600). We therefore also quantify entrainment with the pairwise phase consistency (PPC) [32],

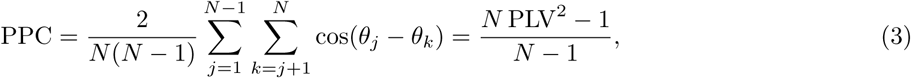

where the equality holds when PLV and PPC are computed from the same N spike phases. For independent phase samples, PPC is an unbiased estimator of the squared population phase-locking value; for large N, PPC *≈* PLV^2^. Sample PPC values can be negative, and PPC is undefined for N < 2.

#### 4.5.2 Statistical testing

Changes in neurons’ preferred phases between conditions were assessed using Moore’s paired-sample circular test [60, 61], applied to paired preferred-phase measurements from individual cells. These comparisons assessed the phase response; locking strength was characterized separately using PLV and PPC. Differences in network response relative to baseline were tested using a fixed-sequence (hierarchical) procedure [62]. For each combi- nation of network state and stimulus frequency, amplitudes were tested in a prespecified order, from highest to lowest, under the assumption that the effect does not decrease with increasing field strength. Testing stopped at the first nonsignificant comparison, and the lowest significant amplitude was reported as the lowest tested amplitude with a detectable change in phase response. The familywise error rate (FWER) was allocated equally across the k dose-response curves using a Bonferroni correction (α_adjusted_ = α/k), controlling the FWER at α = 0.05 under arbitrary dependence between tests. There were five dose-response curves (k = 5), one for each combination of network state and stimulus frequency analyzed. These analyses were performed for all cells combined and separately for the pyramidal cell and inhibitory interneuron populations. The same fixed- sequence procedure was used to test differences in network response between the two network states at each stimulus frequency. These comparisons comprised two dose-response curves (k = 2), one at 7 Hz and one at 10 Hz. Reported p-values were adjusted for multiple comparisons using the Bonferroni correction described above. Because the large number of neurons allows even small shifts to reach statistical significance, effect sizes (differences in preferred phase and in PLV or PPC) are reported alongside p-values.

### 4.6 Implementation and code availability

#### 4.6.1 Implementation

Simulations are performed using the Neurodamus [58] simulation control application, with the NEURON com- putation engine for simulations with extracellular stimulation and the CoreNEURON computation engine for baseline simulations; the two engines to produce identical results by design [63]. The simulations of the single- column circuit reported in the main text were run on a machine whose specifications are shown in Table 2. Baseline simulations required 13 hours of execution time to simulate 5 seconds of biological time, while simu- lations with extracellular stimulation required 35 hours of execution time to simulate 2.5 seconds of biological time. The seven-column circuit simulations and the corresponding single-column circuit simulations reported in Supplementary Section S1 were run on the OBI platform. All simulations and circuits are specified in the SONATA [64] format.

**Table 2:** Computational specifications of the workstation used to run the simulations.

| **Component** | **Specification** |
|---|---|
| System architecture | 64-bit, x86-64 |
| Processor | AMD EPYC 9554 |
| CPU cores | 64 cores |
| CPU clock speed | 3.1 GHz |
| Installed memory | 1.267 TB |
| Storage capacity | 25.18 TB |
| Graphics processor | No dedicated GPU |

#### 4.6.2 Code availability

- The seven-column subvolume of the BBP circuit model is available under the following DOI: https://doi.org/10.5281/zenodo.7930276.
- Neurodamus is available at https://github.com/openbraininstitute/neurodamus
- Code for running the simulations and generating the figures in this paper is available at https://github.com/ITISFoundation/tACS-biophysical-modeling.

#### 4.6.3 Declaration of generative AI and AI-assisted technologies in the writing process

Claude (Sonnet 5 and 5.5, Opus 4) was used to generate code for visualization of preferred spike phases, to generate Fig. 1c-h, and to format Supplementary Table S.1. Claude Sonnet 5 and ChatGPT 5 were used to generate BibTeX citations. Claude and ChatGPT were used to review the manuscript for internal consistency and citation accuracy, to suggest links to biological conditions, and to suggest edits to phrasing for clarity and conciseness. All AI outputs were manually reviewed by the authors.

## Acknowledgments

This research was supported in part by the Swiss National Science Foundation (SNSF) through the Weave/Lead Agency project “Minimally Invasive Targeting of Deep Brain Structures to Treat Epilepsy Using Temporal Interference Stimulation” (grant number 10003470), by a grant from the Swiss National Supercomputing Centre (CSCS) under project ID s1332 on Alps, and by an allocation of computational resources from the Open Brain Institute. JT was supported by a Katja Pokovíc Research Project grant. We thank Alessandro Fasse and Victor Garvalov for technical assistance.

## Author contributions

EH: Methodology, Software, Investigation, Formal Analysis, Visualization, Data Curation, Validation, Writing - original draft, Writing - review and editing JT: Conceptualization, Methodology, Software, Investigation, Formal Analysis, Visualization, Validation, Writ- ing - original draft, Writing - review and editing, Project Administration, Supervision WJ: Software, Validation JK: Software, Validation JBI: Project Administration, Data Curation, Validation MWR: Project Administration, Resources GK: Resources NK: Resources TN: Conceptualization, Writing - review and editing, Supervision EN: Conceptualization, Methodology, Formal Analysis, Writing - review and editing, Supervision

## Supplementary material

### S1 Comparison of single- and seven-column responses

For 10 Hz stimulation in the REM-like network state (see Sec. 2.2), we compare the response of a single cortical column (approximately 30,000 neurons) to a seven-column circuit (approximately 210,000 neurons). The peak- normalized population firing rates show that, after the initial transient, the seven-column circuit has higher relative activity than the single-column circuit during both the baseline period and the stimulated period (Fig. S.1a). At stimulation amplitudes of 0.25 V m*^−^*^1^ and 1 V m*^−^*^1^, the phase preferences of spikes in the two circuits (Fig. S.1c-d) do not differ significantly (Kuiper’s test, p = 0.5 and p = 0.16, Bonferroni-adjusted significance threshold (α*^∗^* = 0.0167)). For a stimulation amplitude of 4 V m*^−^*^1^, the spike phase histograms of the two circuits (Fig. S.1e) do differ significantly (Kuiper’s test, p = 0.008,Bonferroni-adjusted significance threshold (α*^∗^* = 0.0167)), but the difference is on the order of only 10 degrees, and the pairwise phase consistency values are nearly identical (Fig. S.1b). The two circuit sizes support similar entrainment magnitudes, and only small differences in phase preferences.

---

**Figure S1:**
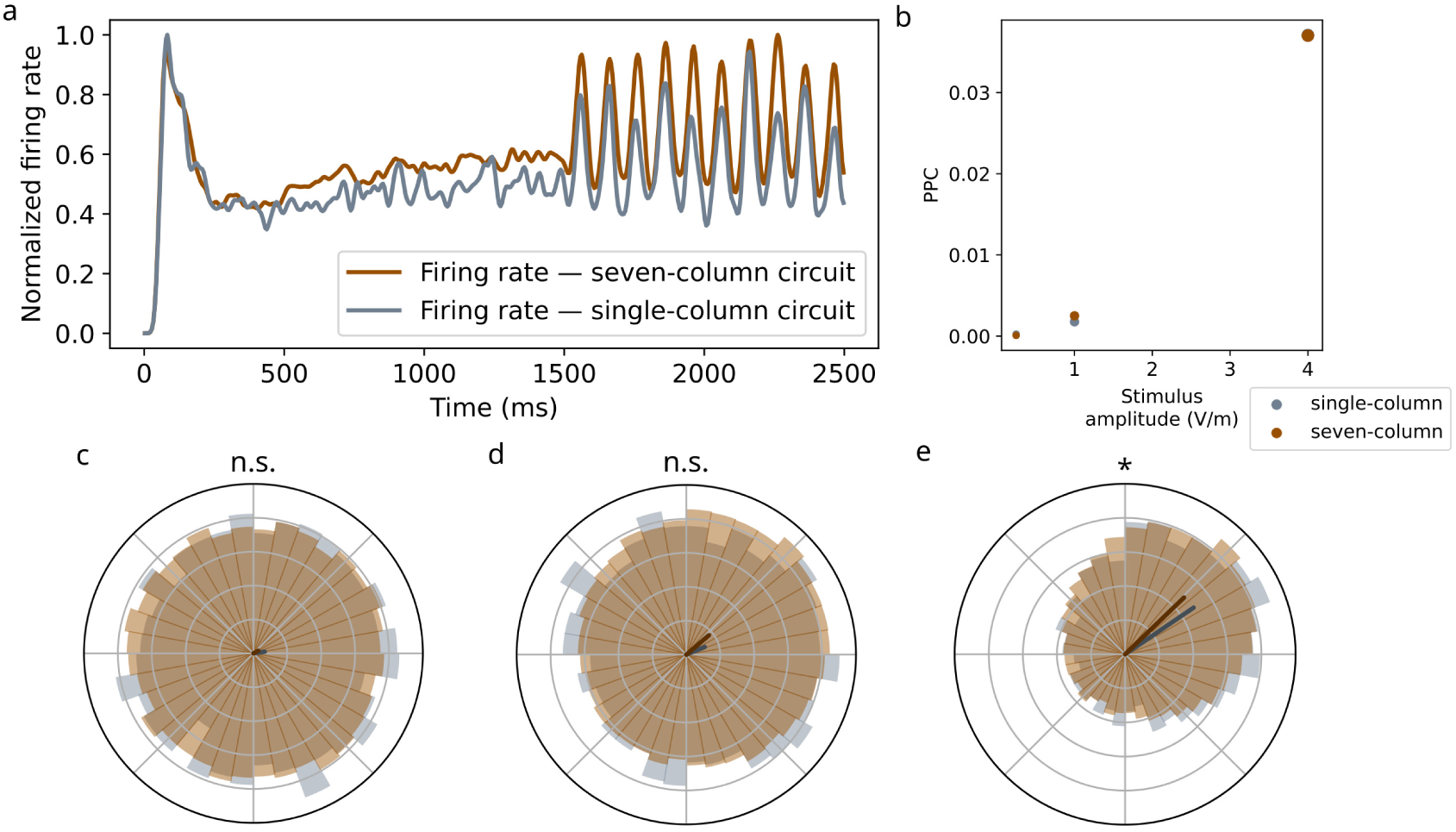
Comparison of single- and seven-column responses at. 10 Hz. a: Smoothed, normalized firing rate histograms of single-column and seven-column circuits, with 10 Hz stimulation at 4 V m*^−^*^1^ applied at t = 1500 ms. b: PPC of the single-column and seven-column circuits in response to 10 Hz stimuli applied at various amplitudes. c-e: Phase distributions for all spikes in the single-column and seven-column circuits in response to 10 Hz stimulus at amplitudes of 0.25 V m*^−^*^1^, 1 V m*^−^*^1^, and 4 V m*^−^*^1^, respectively. Arrow direction indicates mean phase, and length indicates PLV (PPC *≈* PLV^2^ for large spike counts).

**Figure S2:**
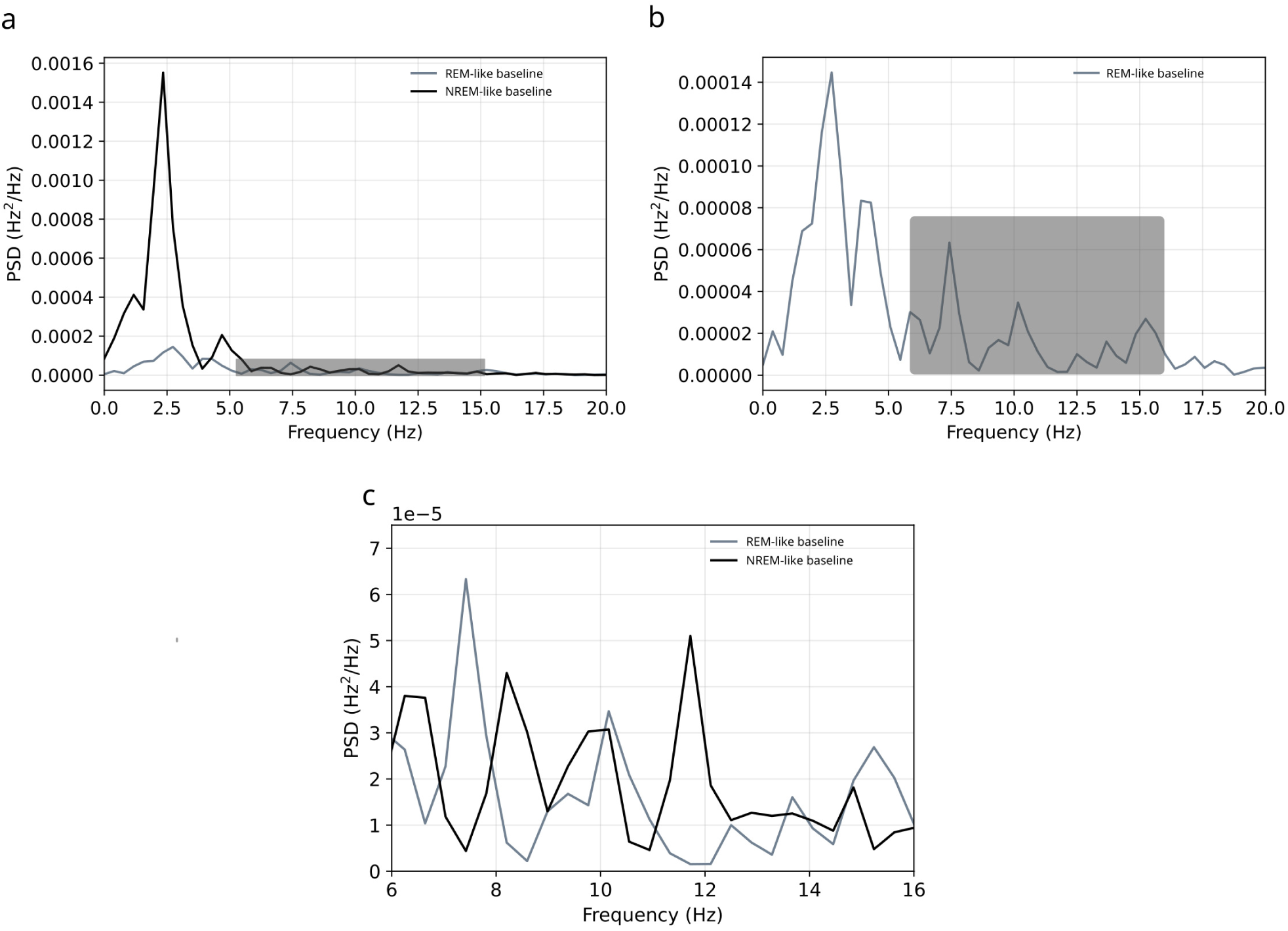
Baseline power spectra of the two network states. a: In the absence of stimulation, the network has no strong intrinsic oscillations in the REM-like network state. In contrast, in the NREM-like state the network exhibits a strong 2.5 Hz oscillation. The gray box indicates the region visualized in panel c. b: In the REM-like state, the network exhibits broadband activity, with the largest peak near 2.5 Hz, and smaller peaks near 7.5 Hz, 10 Hz and 15 Hz. The gray box indicates the region visualized in panel c. c: In the NREM-like state, the peak near 7.5 Hz is absent and peaks appear near 8 Hz and 11.7 Hz, while power near 10 Hz and 15 Hz is slightly lower than in the REM-like state.

**Figure S3:**
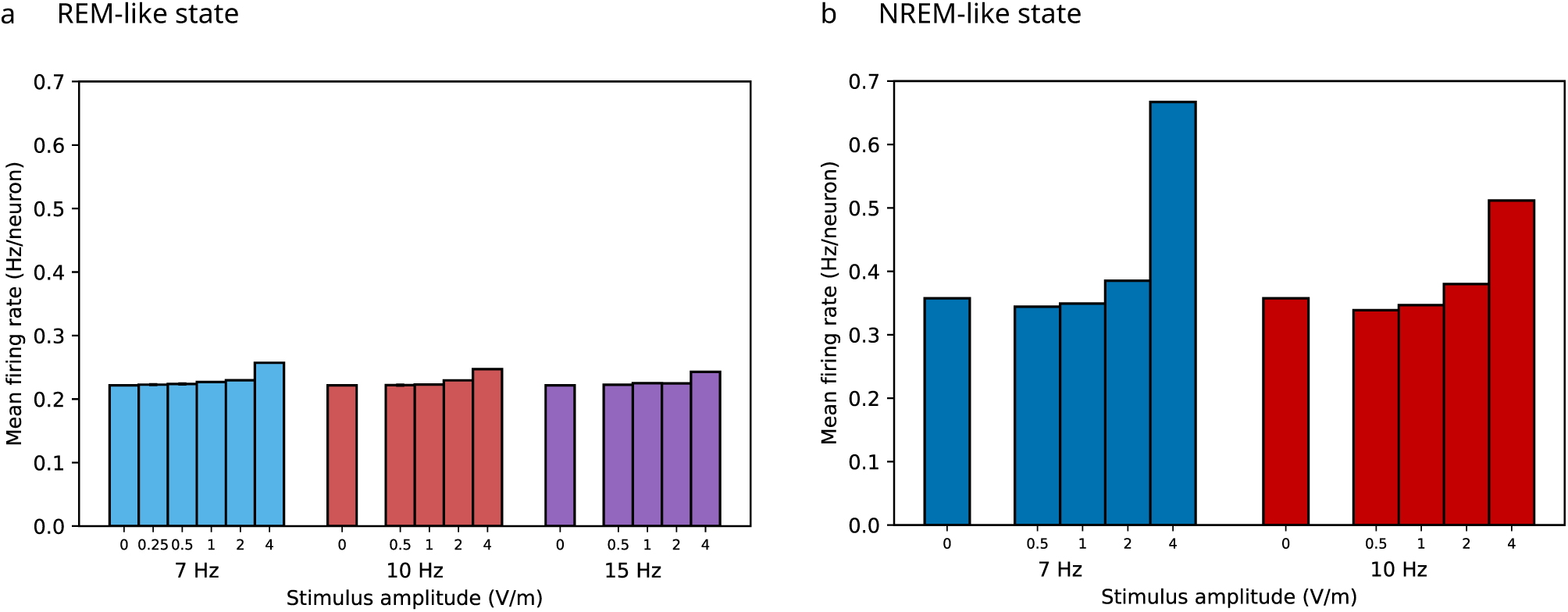
Mean firing rates during stimulation. a: In the REM-like network state, extracellular stimu- lation increases the overall firing rate for all frequencies at an amplitude of 4 V m*^−^*^1^ by approximately 10-20%; for amplitudes 2 V m*^−^*^1^ the increase is less than 5%. b: In the NREM-like network state, the mean firing rate changes little below 2 V m*^−^*^1^ and increases at 4 V m*^−^*^1^ by approximately 40% for 10 Hz and 85% for 7 Hz stimulation. The dashed line indicates the unstimulated baseline.

**Figure S4:**
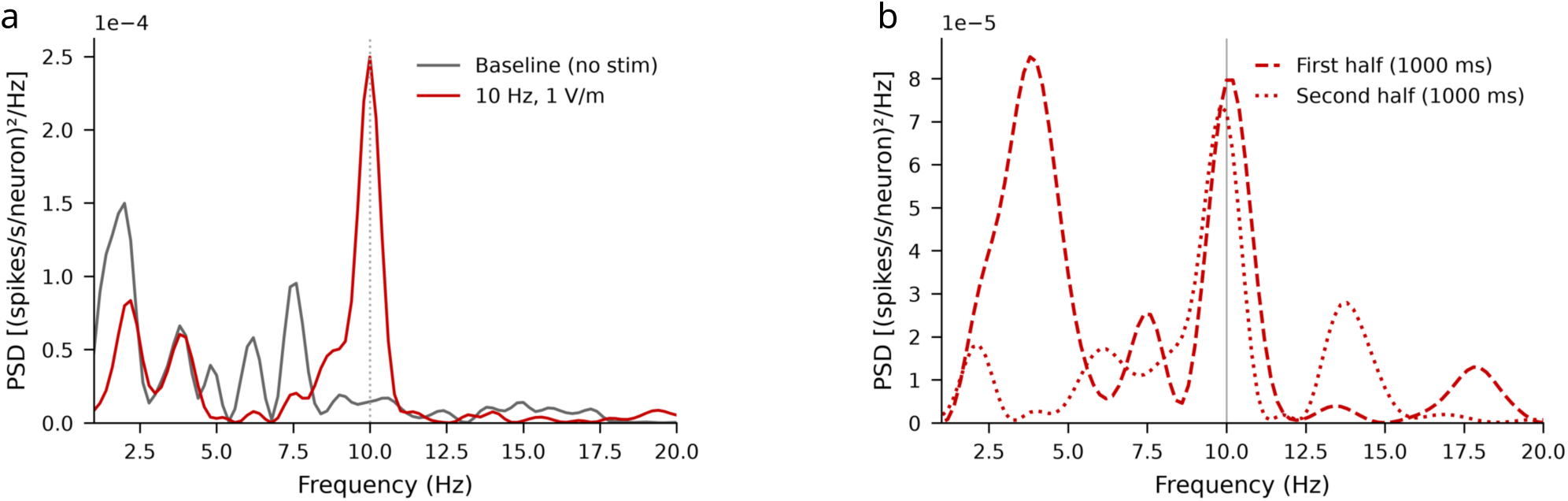
Spectral changes during. 10 Hz **stimulation (REM-like state).** a: During stimulation at 1 V m*^−^*^1^, the PSD of network activity has substantially increased power at the target frequency, and reduced power at off-target frequencies, compared to baseline. b: In the second half of the 2 s stimulation window used in this experiment (as opposed to the 1 s stimulation window used elsewhere), the network exhibits reduced power at most frequencies compared to the first half of the stimulation window.

**Figure S5:**
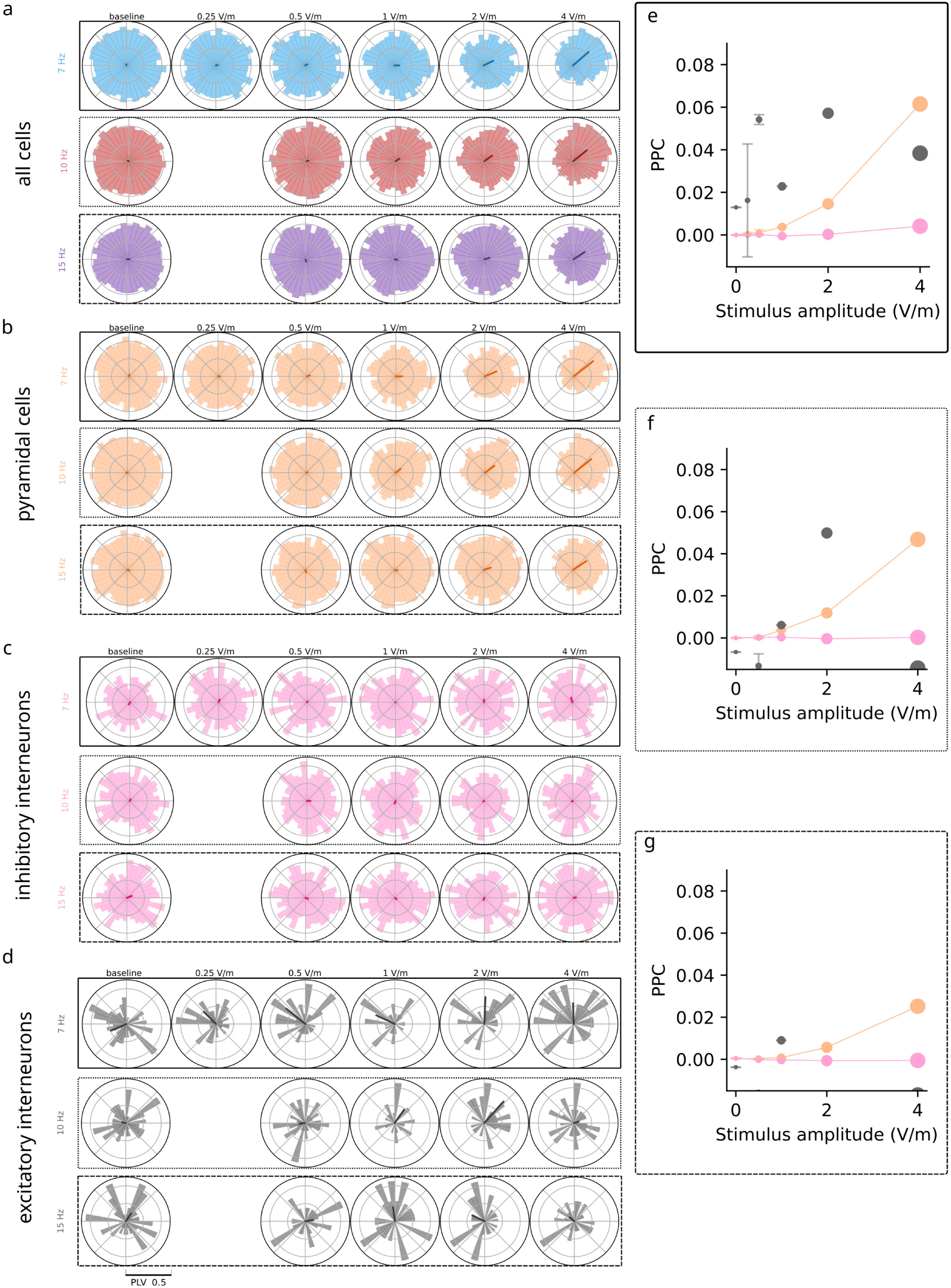
Preferred-phase distributions and PPC by cell type, REM-like state. a: Grid of dis- tributions of cells’ preferred spike phase, from each combination of stimulus amplitude and frequency, in the REM-like network state. b-d: As in a, but for pyramidal cells, inhibitory interneurons, and excitatory interneu- rons, respectively. e-g: PPC of different cell types (colors as in Fig. 2e-h) in response to extracellular stimulus at 7 Hz, 10 Hz, and 15 Hz, respectively. Boxes around each panel correspond to the respective frequencies high- lighted in panels b-d.

**Figure S6:**
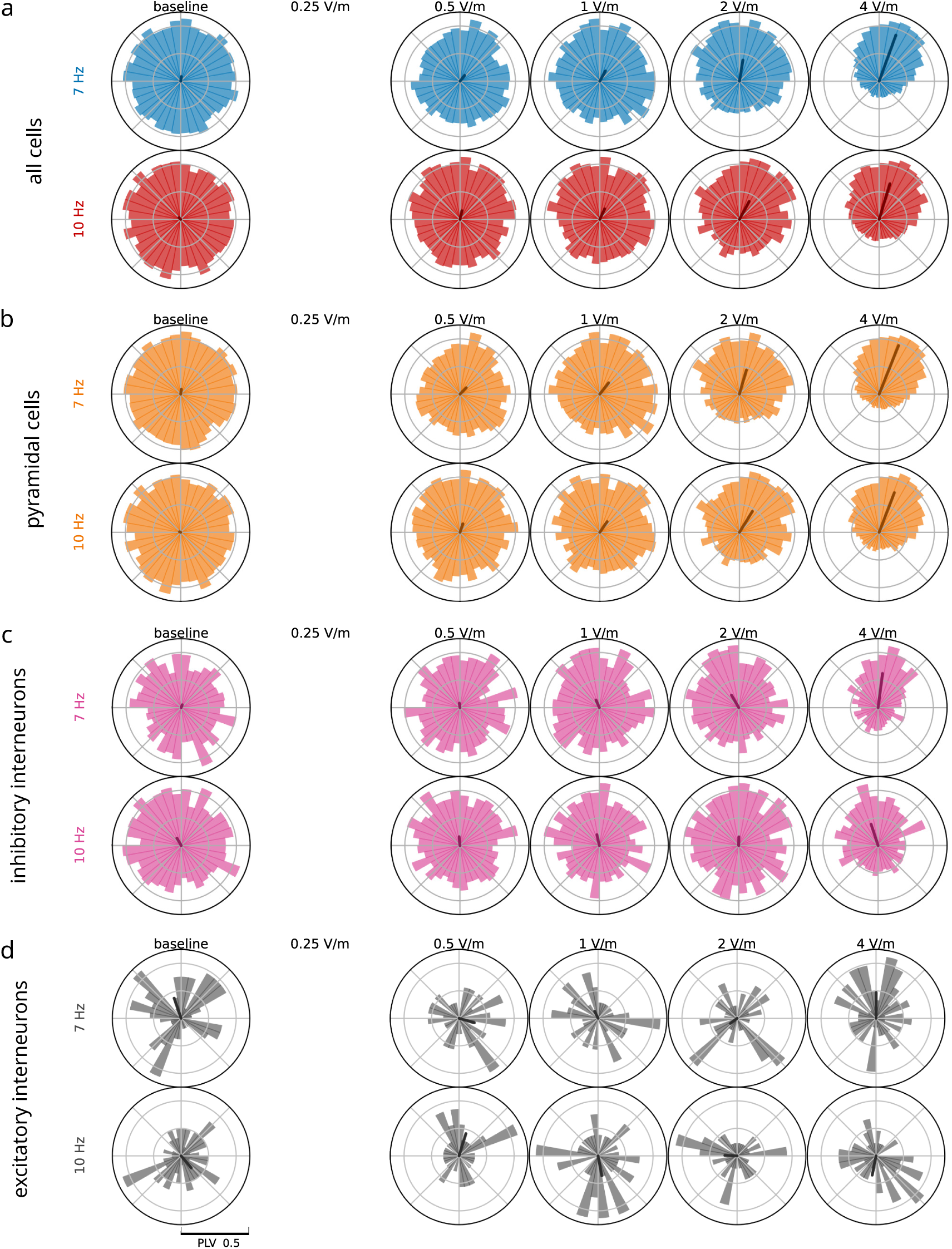
Preferred-phase distributions by cell type, NREM-like state. a-d: As in Supplementary Fig. S.5a-d, but for the NREM-like network state.

**Figure S7:**
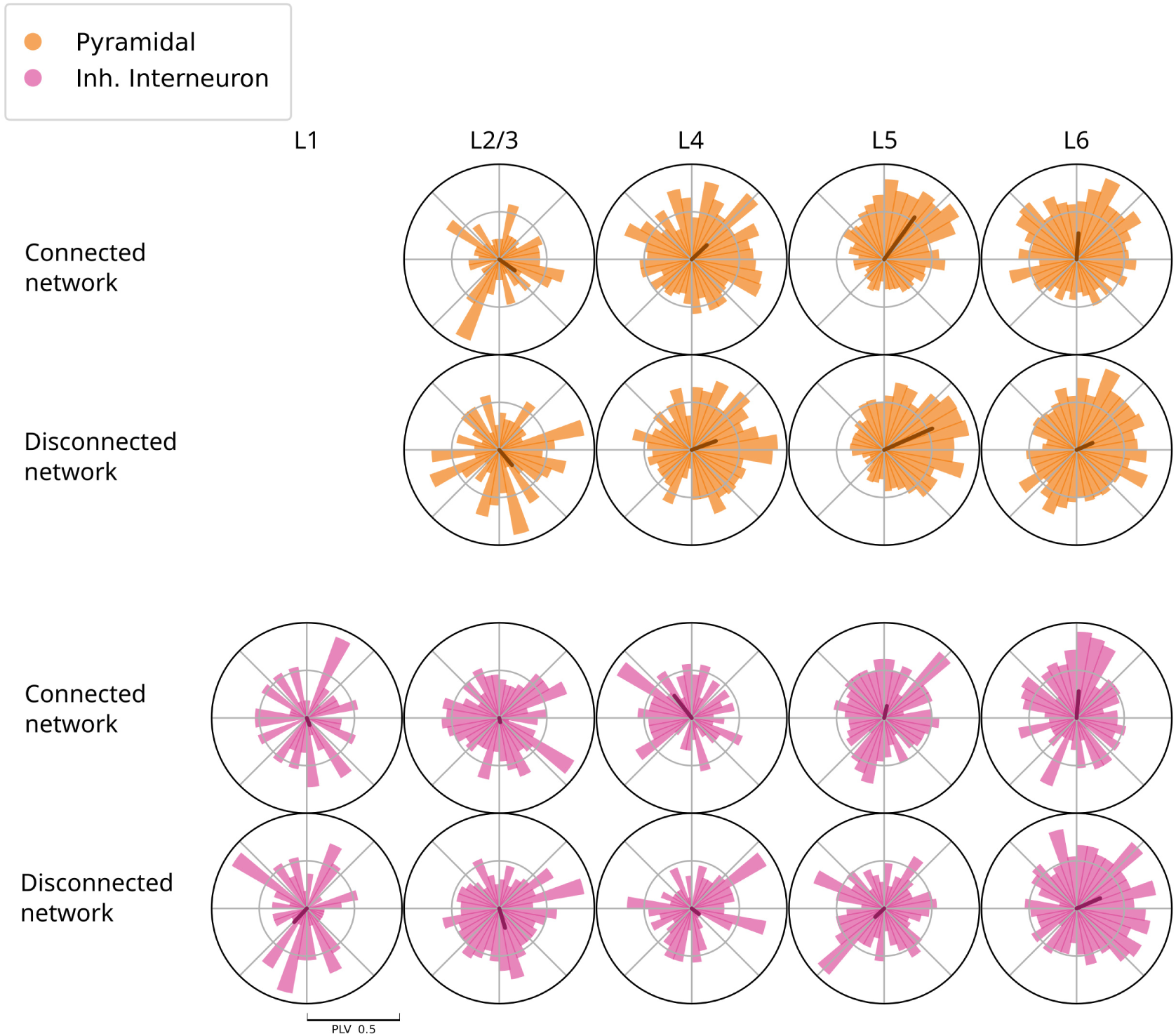
**Connected versus disconnected responses by layer and cell type, NREM-like state (**10 Hz, 2 V m*^−^*^1^). Preferred-phase distributions of pyramidal cells (top) and inhibitory interneurons (bottom) in each layer of the cortical column, for the connected and disconnected (replay) conditions; arrows indicate the resultant vector of each distribution. Differences in preferred phase between the two conditions are largest for layer 5 and layer 6 pyramidal cells and for layer 4 to 6 interneurons, the latter with weak phase concentration. Note that no pyramidal neurons are present in layer 1.

**Table S1:** Mean axonal conduction delay (in milliseconds) between excitatory (E) and inhibitory (I) populations in each layer of the circuit, computed from the connectome.

|  |  |  | **Post** |  |  |  |  |  |  |  |  |  |  |
|---|---|---|---|---|---|---|---|---|---|---|---|---|---|
|  |  |  | L1 | L2 |  | L3 |  | L4 |  | L5 |  | L6 |  |
|  |  |  | I | E | I | E | I | E | I | E | I | E | I |
| **Pre** | L1 | I | 1.1 | 1.2 | 1.2 | 1.2 | 1.4 | 1.3 | 2.6 | 1.2 | 2.5 | 2.1 | 2.7 |
|  | L2 | E | 1.5 | 1.1 | 1.1 | 1.2 | 1.1 | 1.9 | 2.2 | 1.9 | 3.0 | 3.8 | 4.7 |
|  |  | I | 1.2 | 1.2 | 1.2 | 1.1 | 1.3 | 1.5 | 2.0 | 1.3 | 2.9 | 3.0 | 4.3 |
|  | L3 | E | 2.2 | 1.8 | 1.7 | 1.2 | 1.2 | 1.3 | 1.3 | 1.9 | 2.2 | 2.6 | 3.5 |
|  |  | I | 1.5 | 1.3 | 1.2 | 1.1 | 1.1 | 1.3 | 1.4 | 1.4 | 2.1 | 2.5 | 3.5 |
|  | L4 | E | 3.4 | 3.0 | 2.9 | 2.0 | 1.9 | 1.2 | 1.1 | 1.5 | 1.4 | 2.2 | 2.9 |
|  |  | I | 2.7 | 2.2 | 2.0 | 1.2 | 1.3 | 1.1 | 1.1 | 1.4 | 1.4 | 1.5 | 2.5 |
|  | L5 | E | 4.5 | 4.1 | 3.9 | 3.2 | 3.0 | 2.2 | 1.9 | 1.4 | 1.3 | 1.5 | 1.5 |
|  |  | I | 4.0 | 3.6 | 3.5 | 2.2 | 1.9 | 1.3 | 1.3 | 1.3 | 1.2 | 1.4 | 1.6 |
|  | L6 | E | 6.0 | 5.8 | 5.6 | 5.0 | 4.9 | 4.2 | 3.9 | 2.4 | 2.2 | 1.2 | 1.2 |
|  |  | I | 5.6 | 5.3 | 5.1 | 4.0 | 3.6 | 3.2 | 2.8 | 1.7 | 1.5 | 1.1 | 1.2 |

